# A Data Citation Roadmap for Scholarly Data Repositories

**DOI:** 10.1101/097196

**Authors:** Martin Fenner, Mercè Crosas, Jeffrey Grethe, David Kennedy, Henning Hermjakob, Philippe Rocca-Serra, Gustavo Durand, Robin Berjon, Sebastian Karcher, Maryann Martone, Timothy Clark

**Affiliations:** DataCite, Hannover, Germany; Institute for Quantitative Social Science, Harvard University, Cambridge MA, USA; University of California San Diego, La Jolla CA, USA; University of Massachusetts Medical School, Worcester MA, USA; European Bioinformatics Institute (EMBL-EBI), European Molecular Biology Laboratory, Hinxton, Cambridgeshire, UK; Oxford e-Research Centre, University of Oxford, Oxford, UK; Standard Analytics, New York NY, USA; Qualitative Data Repository, Syracuse University, Syracuse NY, USA; Massachusetts General Hospital, Boston MA, USA.; Harvard Medical School, Boston MA, USA.

**Author notes:** These authors contributed equally to this work (Martin Fenner, Mercè Crosas). Corresponding author: Martin Fenner.

## Abstract

This article presents a practical roadmap for scholarly data repositories to implement data citation in accordance with the Joint Declaration of Data Citation Principles, a synopsis and harmonization of the recommendations of major science policy bodies. The roadmap was developed by the Repositories Expert Group, as part of the Data Citation Implementation Pilot (DCIP) project, an initiative of FORCE11.org and the NIH BioCADDIE (https://biocaddie.org) program. The roadmap makes 11 specific recommendations, grouped into three phases of implementation: a) required steps needed to support the Joint Declaration of Data Citation Principles, b) recommended steps that facilitate article/data publication workflows, and c) optional steps that further improve data citation support provided by data repositories.

## Introduction

The Joint Declaration of Data Citation Principles (JDDCP) published in 2014 (Data Citation Synthesis Group, 2014) and endorsed by a large number of scholarly and academic publishing organizations, lays out a set of principles on purpose, function and attributes of data citations, starting with stressing that data should be considered legitimate, citable products of research (M Altman, Borgman, & Crosas, 2015). The JDDCP condense the results of substantial prior studies on science policy and practice (King, Gary & Altman, Micah, 2007; Uhlir, 2012; CODATA-ICSTI Task Group on Data Citation Standards and Practice, 2013).

The JDDCP intentionally focuses on data citation principles, as the implementation of these principles will differ across disciplines and communities. The roadmap presented here aims to provide practical guidance for repositories on implementing these data citation principles with a focus on life sciences, based on earlier work in this area, in particular Starr et al. (2015) and Altman and Crosas (2013), and are consistent with recent recommendations regarding data, code and workflows (Smith, Katz, & Niemeyer, 2016; Stodden et al., 2016).

Data repositories play a central role in data citation, as they provide stewardship and discovery services to find data, give persistent access to the data being cited, and provide unique identifiers and metadata needed for data citation. For data citation, repositories need to work closely with a variety of stakeholders, including publishers, reference manager providers, and of course researchers. Data citation practices and technologies supported by repositories will substantially assist development of new data discovery indexes such as BioCADDIE.

## Results

The guidelines are grouped into three phases: required, recommended and optional. Implementing these guidelines takes time and resources, it is therefore not only critical to provide specific guidelines, but also to give guidance on priorities: work needed to support the Joint Declaration of Data Citation Principles (required phase), additional work to facilitate article/data publishing workflows in collaboration with publishers (recommended phase), and extra work to support data citation that can be done by data repositories (optional phase). The Guidelines are summarized in Table 1, and discussed in detail in the text following the table.

**Table 1.** Guidelines for Repositories.

| Level | # | Guideline |
| --- | --- | --- |
| Required | 1 | All datasets intended for citation *must* have a globally unique persistent identifier that can be expressed as unambiguous URL. |
|  | 2 | Persistent identifiers for datasets *must* support multiple levels of granularity, where appropriate. |
|  | 3 | This persistent identifier expressed as URL *must* resolve to a landing page specific for that dataset. |
|  | 4 | The persistent identifier *must* be embedded in the landing page in machine-readable format. |
|  | 5 | The repository must provide documentation and support for data citation. |
| Recommended | 6 | The landing page *should* include metadata required for citation, and ideally also metadata helping with discovery, in human-readable and machine-readable format. |
|  | 7 | The machine-readable metadata *should* use schema.org markup in JSON-LD format. |
|  | 8 | Metadata *should* be made available via HTML meta tags to facilitate use by reference managers. |
|  | 9 | Metadata *should* be made available for download in Bibtex and/or another standard bibliographic format. |
| Optional | 10 | Content negotiation for schema.org/JSON-LD and other content types *may* be supported so that the persistent identifier expressed as URL resolves directly to machine-readable metadata. |
|  | 11 | HTTP link headers *may* be supported to advertise content negotiation options |

Details of each recommendation follow, with examples.

### 1. Persistent identifiers

A data citation *must* include a persistent method for identification that is machine actionable, globally unique, and widely used by a community (**JDDCP**, principle #4). For implementation by data repositories, this means:

- **Persistent method for identification.** Unique identifiers, and metadata describing the data, and its disposition, *must* persist -- even beyond the lifespan of the data they describe (**JDDCP**, principle #6). As extension to this principle data repositories should make provisions to keep unique identifiers and metadata available beyond the lifespan of the data or repository, ideally in a well-recognized and accepted standard metadata format.
- **Machine actionable**. The persistent identifier *must* be understood, and be resolvable, as an HTTP URI in accordance with the RFC 3986 (“RFC 3986 - Uniform Resource Identifier (URI): Generic Syntax,” 2005), including support for content negotiation (Treloar, 2011).
- **Globally unique.** The identifier *must* use a prefix (namespace) if the identifier character string is only unique within a particular database, e.g. an accession number. For data repositories that are not using globally unique identifiers, the DCIP EG2 Identifiers Expert Group_is working on a bridging solution using common prefixes (“Prefix Commons,” 2016) and resolver services (identifiers.org and n2t.net).
- **Widely used by a community.** Accession numbers, in combination with the database name for global uniqueness, are the most widely used identifiers in the life sciences.

### 2. Persistent identifier granularity

Persistent identifiers for datasets must support multiple levels of granularity to support both the citation of a specific version and/or individual dataset, as well the citation of an unspecified version of a dataset and/or a collection of primary data.

In many domains, primary data is uniquely identified and cited as a collection of potentially many individual items. At the same time, these individual items need their own unique identifiers to support later reuse and recombination into different sets while maintaining the ability to cite the constituent data elements.

An example is in the field of neuroimaging, where individual subject scans using a given imaging modality are the lowest level at which objects will be identified, while the primary publication will cite a collection level unique identifier. This imposes a requirement that lower-level identifiers need to be able to be grouped via a collection identifier and accessed as set elements from the overall collection landing page (Honor, Haselgrove, Frazier, & Kennedy, 2016). Another example is the BioStudies database (McEntyre, Sarkans, & Brazma, 2015), which can provide storage for all the underlying data links and files for a publication.

Only in circumstances where multiple levels do not inherently exist in the data, i.e. no collections or other groupings exist, may this requirement be waived.

### 3. Landing pages

The persistent identifier expressed as HTTP URL *must* resolve to a specific landing page for that dataset or dataset collection. The persistent identifier expressed as HTTP URL *must not* resolve to the data itself (Starr et al., 2015), or to other representations of the metadata, unless special protocols such as content negotiation are used (see guideline 7 below).

**Figure 1.**
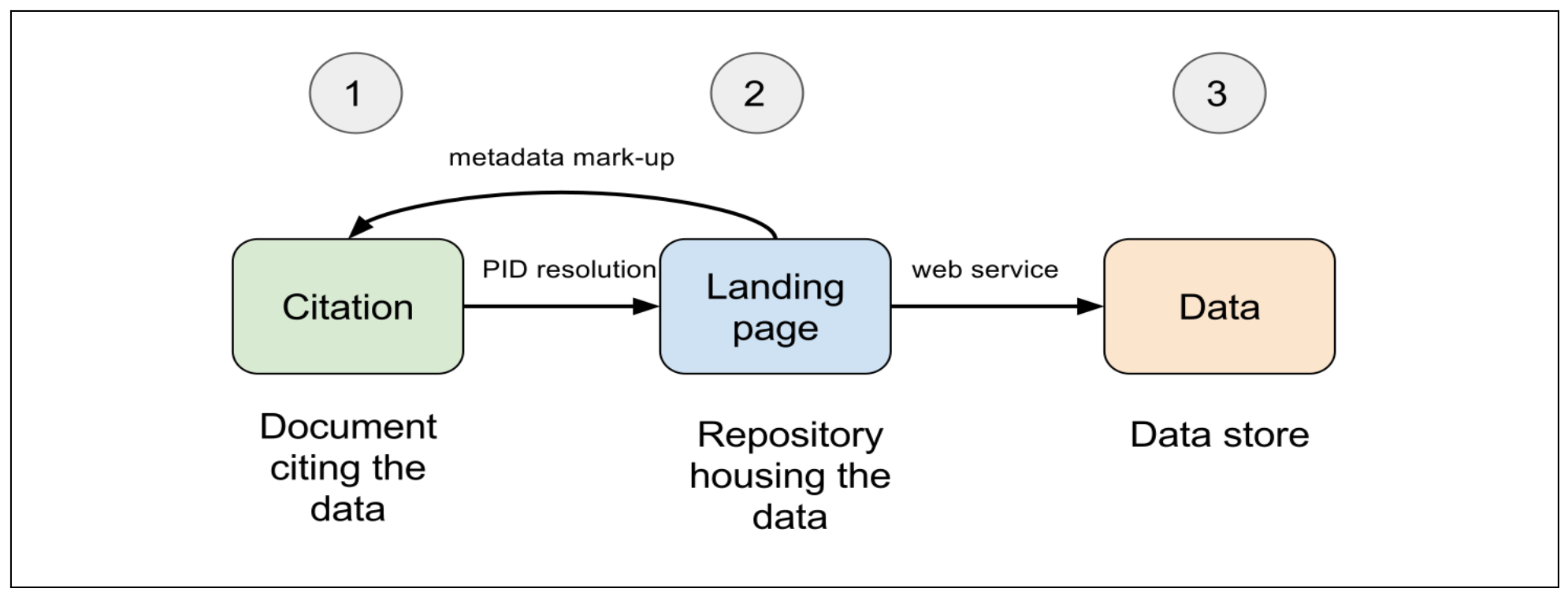
Generic data citation - relationships of the citation reference, repository landing page and underlying data, taken with permission from (“Data Citations: A Primer,” 2016).

Landing pages provide definitive information (metadata) on how the dataset should be cited, other descriptive information about the dataset, as well as data accessibility and licensing information. Repositories should provide a landing page for every dataset or collection of datasets intended to be cited, which could be single entries, sets of entries, the entire repository or a curated database (Starr et al., 2015).

Reference to a statement describing the data and metadata persistence policies of the repository should also be provided at the landing page. Data persistence policies will vary by repository but should be clearly described (Starr et al., 2015). Figure 2 provides an example.

**Figure 2.**
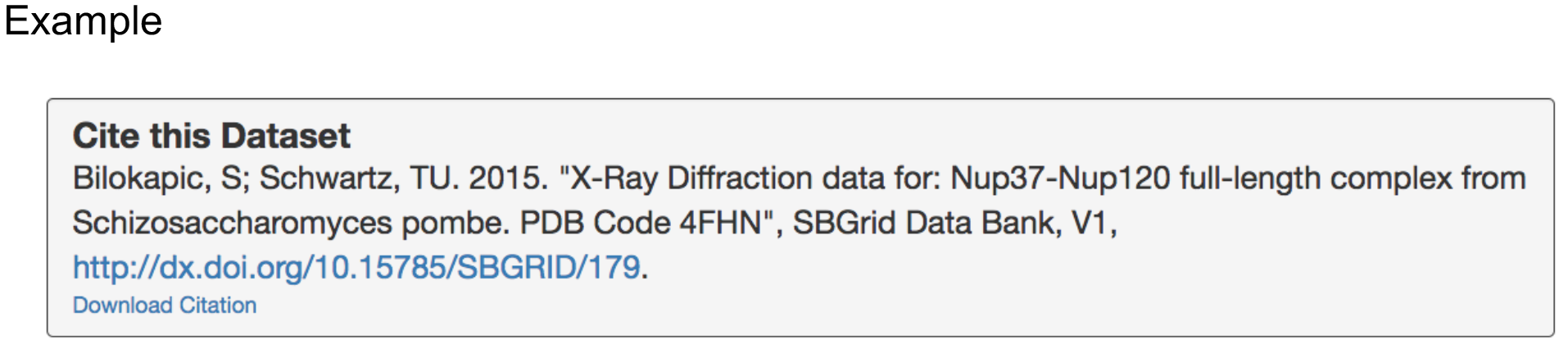
Providing information about how a dataset should be cited, with Bibtex download link.

### 4. Persistent identifiers on landing pages

To verify that a persistent identifier resolves to a correct landing page, the persistent identifier *must* be embedded in the landing page in human-readable and machine-readable formats. This enables basic data citation by reference managers, and enables minimal validation by the publisher of persistent identifiers cited in documents. The persistent identifier should be found somewhere on the landing page, but is ideally embedded in schema.org markup and/or using HTML meta tags.

#### Example schema.org/JSON-LD

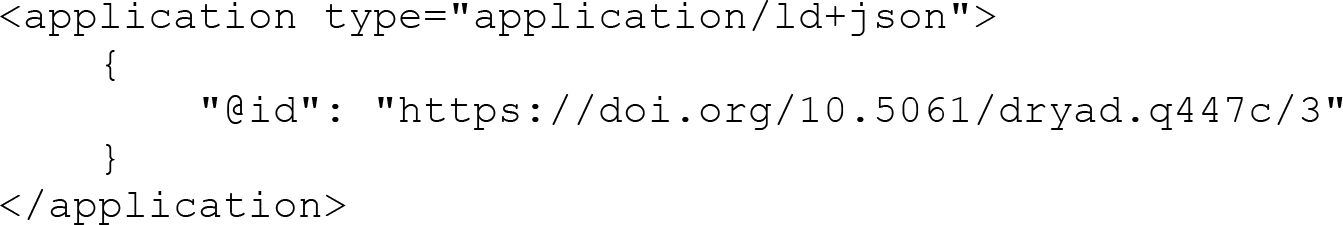

#### Example HTML meta tags

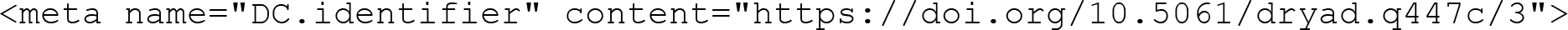

### 5. Documentation and author support

The repository *must* provide documentation about how data should be cited, how metadata can be obtained, and who to contact for more information. The DCIP FAQ Expert Group provides example documentation for data repositories.

### 6. Metadata on landing pages

Landing pages *should* provide metadata required for data citation in both human- and machine-readable format. The latter includes that access to the metadata should not require Javascript, cookies or login. The landing page *should* show the citation metadata in human-readable form, e.g. formatted in one or more citation styles common to the community in a `Cite this Dataset` field and, possibly, provide means of copying/downloading the citation as text. The landing page should also show all versions, or link to a page with version information. A visible link to machine-readable metadata should be provided.

The metadata elements needed for data citation are given in Table 2.

**Table 2:** Citation metadata for Data Repositories.

| Citation Metadata | Dublin Core<sup>a</sup> | Schema.org<sup>b</sup> | DataCite<sup>c</sup> | DATS<sup>d</sup> |
| --- | --- | --- | --- | --- |
| Dataset Identifier | identifier | @id* | identifier | identifier |
| Title | title | name | title | title |
| Creator** | creator | author | creator | creator |
| Data repository or archive | publisher | publisher | publisher | publisher |
| Publication Date | date | datePublished | publicationYear | date |
| Version | *not available* | version | version | version |
| Type | type | type | resourceTypeGeneral | type |

<sup>a</sup> Dublin Core Metadata Element Set (“Dublin Core Metadata Element Set, Version 1.1,” 2012);
<sup>b</sup> Dataset - Schema.org (“Dataset - schema.org,” n.d.);
<sup>c</sup> DataCite Metadata Working Group (2016);
<sup>d</sup> Gonzalez-Beltran & Rocca-Serra (2016);
\* name of ID field depends on schema.org serialization format, it is **@id** for JSON-LD;
\*\* not all datasets will have “the main researchers involved in producing the data” (DataCite Schema), in which case the more generic “An entity primarily responsible for making the resource” from Dublin Core should be used, and this can also be an organization.

All metadata fields required for citation are part of Dublin Core (with the exception of *version*), the core schema.org specification, and by extension Bioschemas (“BioSchemas,” 2016), as well as the DataCite and DATS metadata schemata.

In addition to the metadata required for citation, it is recommended to provide additional metadata on landing pages – again in human-readable and machine-readable formats – that help with data discovery, as shown in Table 3.

The metadata standards Dublin Core, schema.org and DataCite by their very nature of being generic only provide some metadata helpful for discovery, while DATS can provide much more detailed information about a biomedical dataset. Further information can be found in the DataMed DATS specification.

**Table 3:** Important discovery metadata for Data Repositories.

| Discovery Metadata | Dublin Core | Schema.org | DataCite | DATS |
| --- | --- | --- | --- | --- |
| Description | description | description | description | dataType<br>dimension<br>Material<br>...* |
| Keywords | subject | keywords | subject | keywords |
| License | license | license | rights | license |
| Related Dataset** | isPartOf<br>isVersionOf<br>references | isPartOf<br>citation | relatedIdentifier | isPartOf |
| Related Publication*** | bibliographicCitation | citation | relatedIdentifier | publication |

\* DATS provides much more detailed metadata to describe a biomedical dataset;
\*\* related datasets can have part/whole relations (IsPartOf, etc.), version relations (IsVersionOf, etc.) or reference relations (references);
\*\*\* related publications reference a dataset published previously, reference a dataset published in parallel with the publication, or otherwise document a dataset

Information about related datasets should be provided where possible, as should information about related publications. They provide important information that can help with discovery. When a data repository knows about a publication citing a dataset, this information should be included in the metadata, complementing the information about the dataset found in the citing publication and enabling navigation between publication and dataset in both directions.

### 7. Metadata on landing pages using schema.org/JSON-LD

All dataset landing pages *should* provide machine-readable metadata using schema.org markup in JSON-LD format. JSON-LD is the easiest way to represent schema.org metadata, and is also used to represent DATS metadata in schema.org format (Gonzalez-Beltran & Rocca-Serra, 2016). The JSON-LD should be embedded in the HTML page using a `<script type="application/ld+json"` tag.

#### Examples

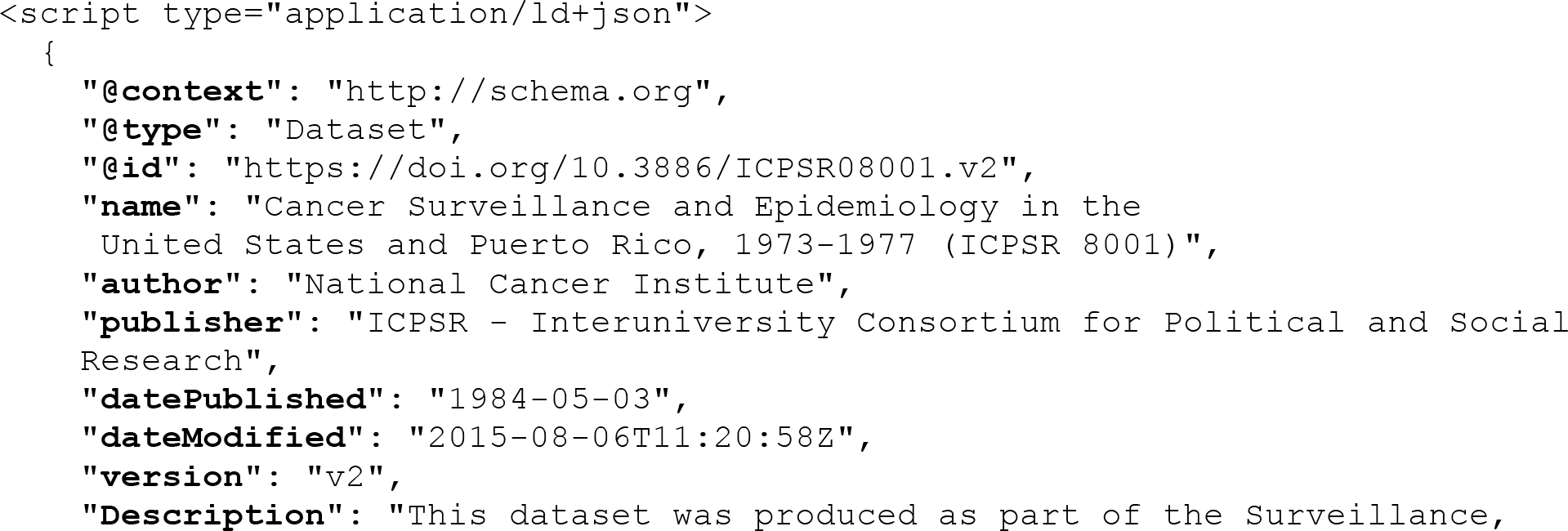

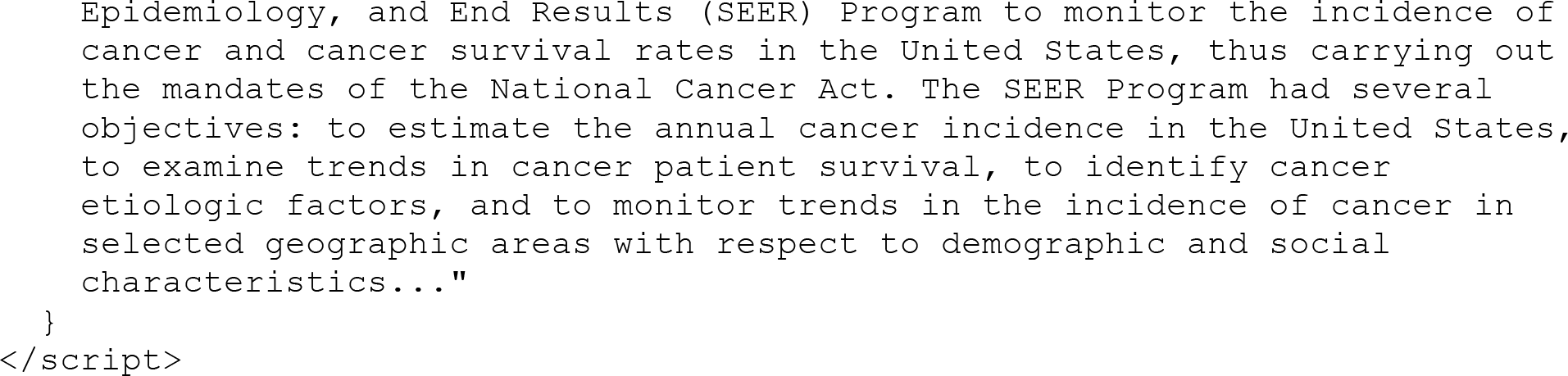

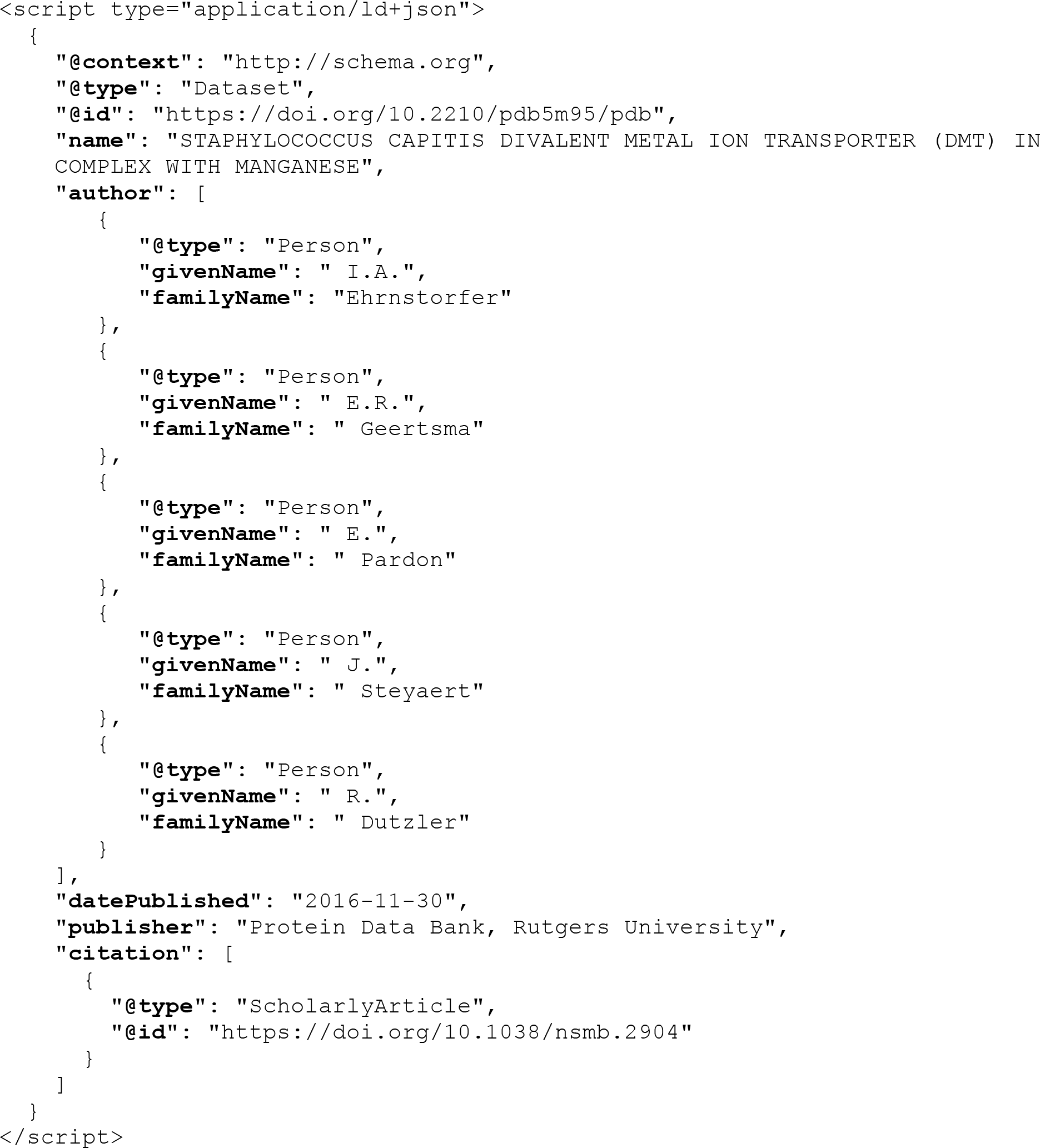

For further examples please use DataCite Search_(“DataCite Search,” 2016), which has embedded schema.org/JSON-LD metadata on every search result page for more than three million datasets.

### 8. Metadata via HTML Meta Tags

Data repositories *should* offer machine-readable metadata on landing pages using Highwire, PRISM (Hammond, Hannay, & Lund, 2004), and/or Dublin Core HTML meta tags. These HTML meta tags are currently the preferred method of reference managers to extract the persistent identifier or full citation metadata from landing pages, as reference managers currently don’t routinely support schema.org/JSON-LD metadata extraction.

#### Example

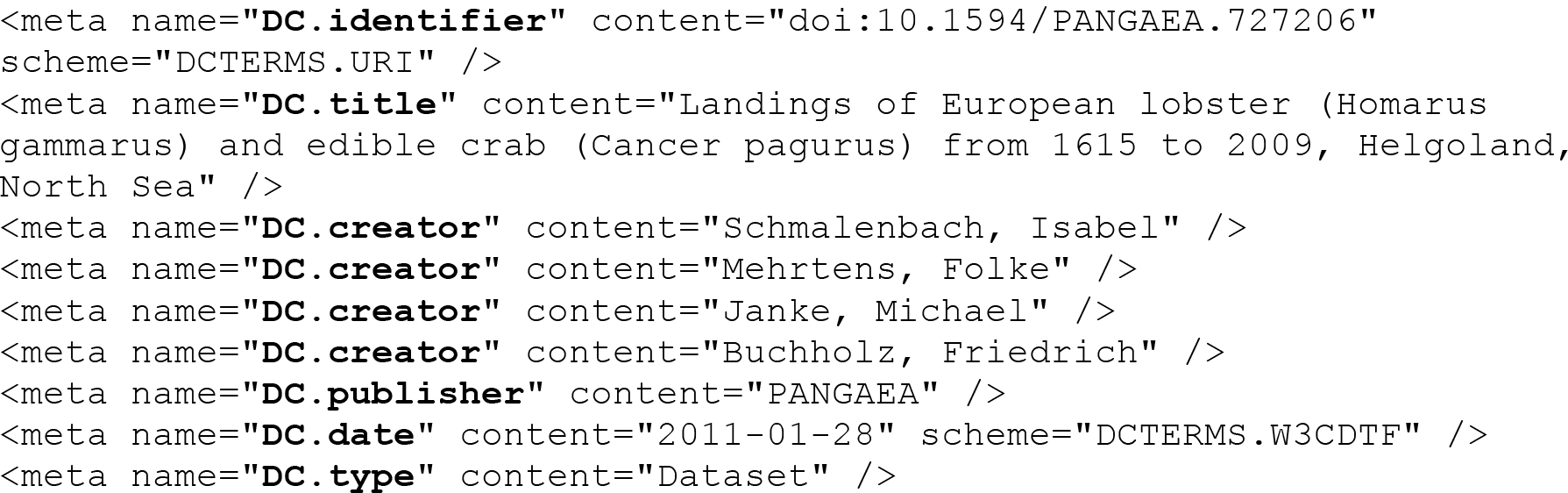

### 9. Metadata via downloadable file in standard bibliographic format

Repositories *should* provide a download link in a common bibliographic format – e.g. .bib (BibTex file format) and/or .ris (RIS file format) – on the landing page of the dataset. The file should include all metadata required for a data citation.

#### Example: BibTex

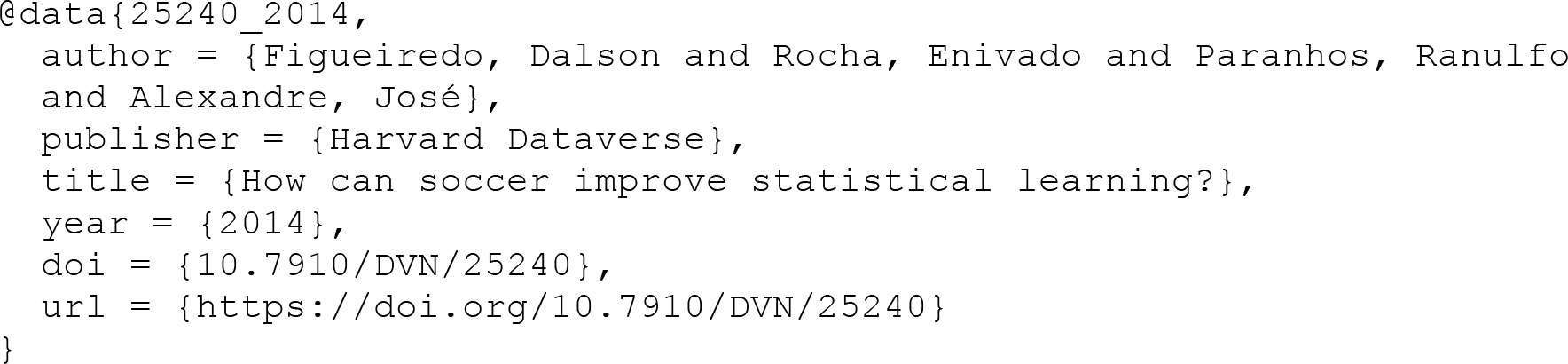

#### Example: RIS

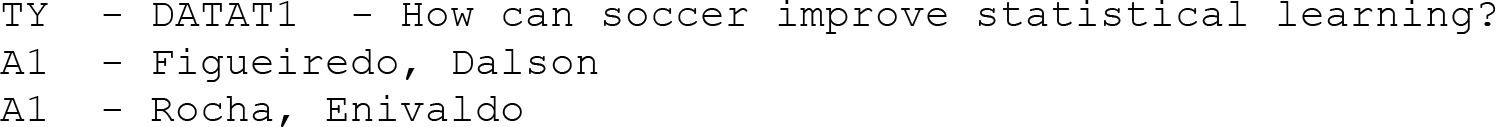

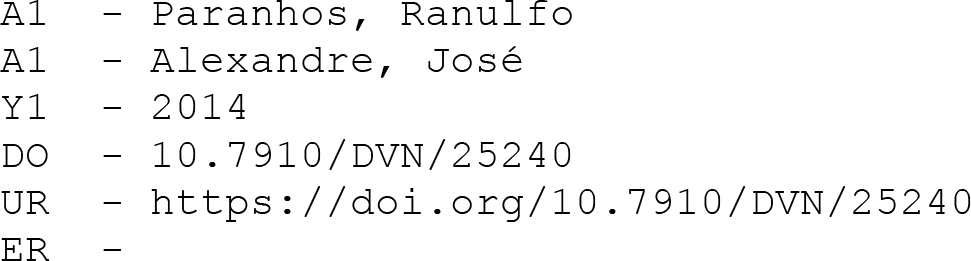

### 10. Content negotiation for machine-readable metadata

Persistent identifiers expressed as HTTP URI *must* by default resolve to the landing page for that dataset (see guideline #3). Data repositories and identifier service providers such as identifiers.org or DataCite in addition *may* implement content negotiation for the persistent identifier expressed as HTTP URI, returning machine readable metadata in various formats. Content negotiation is for example supported by identifiers.org and DataCite and can return metadata in XML, RDF, Bibtex and other metadata formats.

#### Example: Image Attribution Framework (IAF)

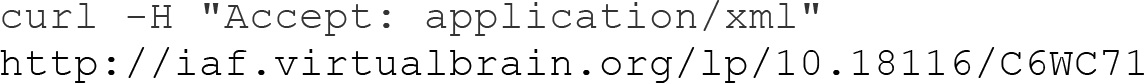

In addition, the HTML version of this page has a link to the XML (available without content negotiation at http://iaf.virtualbrain.org/lp/xml/10.18116/C6WC71).

#### Examples: DataCite

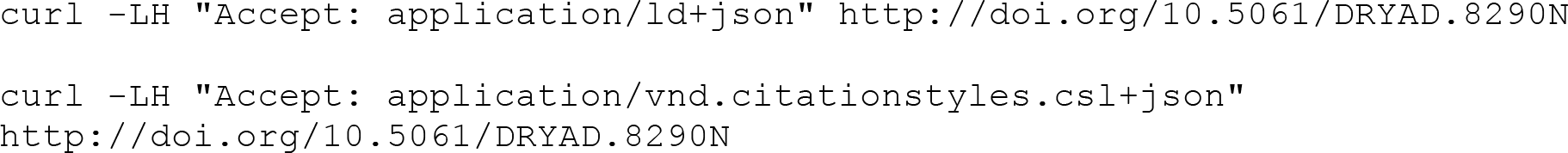

Metadata in `application/vnd.citationstyles.csl+json` format are used as input by many reference managers, e.g. Zotero or Mendeley.

### 11. Support HTTP link headers

The persistent identifier (see guideline #2) and available content negotiation options (see guideline #9) *may* be provided in a HTTP link header (Van de Sompel & Nelson, 2015). This facilitates discovery of content negotiation options and makes it easier to fetch the identifier from large landing pages, as only a HTTP head request is needed).

#### Example

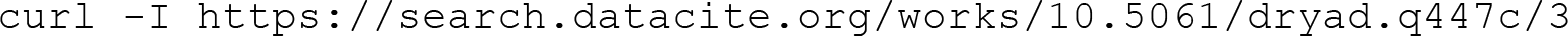

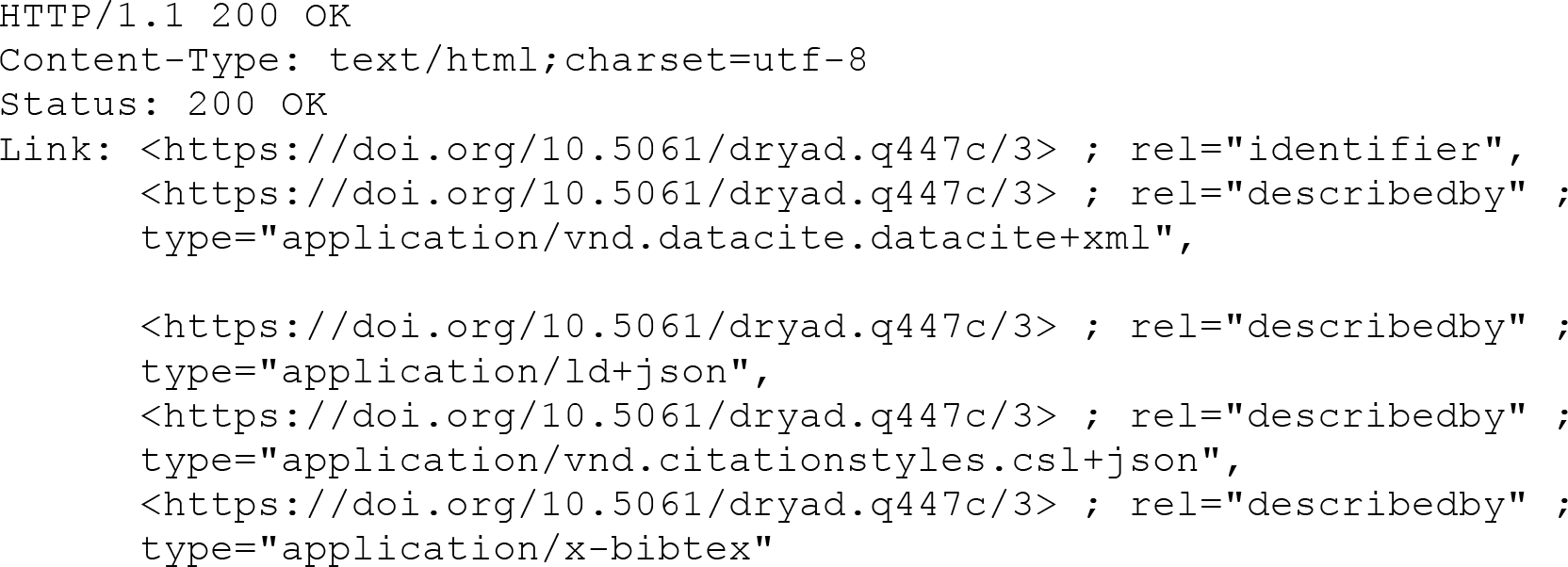

## Methods

This roadmap was developed based on numerous discussions of the DCIP Repositories Early Adopters Expert Group, led by Martin Fenner and Mercè Crosas, including two in-person workshops in February (Boston) and June (San Diego) 2016, and in close coordination with the other DCIP expert groups. The resulting guidelines have been widely circulated. A course on the guidelines and how to implement them, was held at the FORCE11 Scholarly Communication Institute (FSCI) in August of 2017. The course instructors were Martin Fenner and Gustavo Durand, with guest speaker Natasha Noy from the schema.org initiative.

At the conclusion of the course, a hackathon was coordinated by Fenner and Durand, with Noy helping in schema.org metadata integrations. This hackathon was open to the course participants as well as other interested attendees at FSCI. Small teams that included staff from several data repositories were formed and each worked on implementing at least one of the 10 guidelines for their respective data repositories. Overall, the hackathon focused on machine-readable metadata in landing pages, specifically in schema.org JSON-LD, and some repositories had implemented schema.org support by the end of the hackathon.

The course and hackathon provided valuable feedback regarding the guidelines, e.g. that providing metadata in bibtex format for download should be recommended not optional. It served as both a propagation mechanism for the guidelines and a means of validation with practitioners.

An informal poll was taken of course participants as part of this validation. They represented a self-selected sample of technologists working for various biomedical data repositories and well-positioned to implement the guidelines. Results are shown in Table 4. Based on this limited survey, we see that most repositories we queried have implemented, or are in the process of implementing, the required guidelines. Also, of the recommended and optional guidelines, the surveyed attendees expressed most interest in implementing machine-readable metadata in landing pages and in link headers. The guidelines’ overall approach was strongly endorsed.

We expect based on our survey, that many data repositories already follow all required recommendations, but most need to do more work for those that are recommended or optional.

**Table 4.** Informal implementation survey results from FSCI course and hackathon. Some responses indicated “partial implementation” – we were conservative and only counted complete implementations.

| **Guideline** | **Already Implemented** | **Planning to Implement** | **Don’t Care** | **No Response** |
| --- | --- | --- | --- | --- |
| **1** | 16 | 0 | 0 | 0 |
| **2** | 10 | 3 | 2 | 1 |
| **3** | 12 | 2 | 1 | 1 |
| **4** | 10 | 4 | 0 | 2 |
| **5** | 13 | 2 | 0 | 1 |
| **6** | 9 | 4 | 0 | 3 |
| **7** | 1 | 8 | 0 | 6 |
| **8** | 5 | 5 | 1 | 5 |
| **9** | 7 | 2 | 0 | 7 |
| **10** | 6 | 4 | 0 | 5 |
| **11** | 1 | 8 | 0 | 7 |

## Conclusions

This document provides a roadmap for scholarly data repositories to implement support for data citation. Most if not all required steps have already been implemented by many data repositories, and little if any work is needed by them to fully support the Joint Declaration of Data Citation Principles. More work is still needed to implement the recommended steps, and in particular support for schema.org/JSON-LD markup for metadata is still at an early stage. Data repositories that have implemented the required and recommended steps might be interested to look into the optional steps for extra data citation support.

The Data Citation Implementation Pilot and this document focus on data citation support in scholarly data repositories. Using persistent identifiers, standard machine-readable metadata and landing pages of course not only supports data citation, but also facilitates data discovery. Data discovery requires more specific metadata than the metadata needed for data citation, and it is facilitated by a central index of all datasets. The NIH BD2K bioCADDIE project, of which the Data Citation Implementation Pilot is a small part, is working on standard metadata for biomedical data with DATS, and on a central index to search a large number of biomedical datasets with DataMed_(“DataMed | bioCADDIE DDI,” 2016). The European ELIXIR_(“ELIXIR Data for life,” 2016) project (life sciences) and DataCite (all disciplines) are also working on standard metadata and a search index for data discovery. Both Elixir and DataCite are closely collaborating with bioCADDIE in these activities.

The data citation roadmap for scholarly data repositories described in this document is an important step towards full data citation support by data repositories. Going forward a lot of work is needed to implement these guidelines, and ongoing coordination amongst data repositories, and with publishers and other important stakeholders will be essential in this activity.

## Acknowledgements

This document was generated by the DCIP Repositories Expert Group with input from data repositories, publishers, persistent identifier providers, reference manager specialists, and other experts on data citation. Implementation of the data citation principles involves many stakeholder groups, and the DCIP project has worked closely with them via several expert groups, and a coordinating steering group.

Research reported in this publication was supported in part by the National Institutes of Health under award number U24HL126127 for the BioCADDIE project; and by the European Molecular Biology Laboratory (EMBL).

The authors gratefully acknowledge the following members of the Data Citation Repositories Expert Group, who participated actively with the authors in workshops and/or telecons to develop this Roadmap: Cecilia Arighi (Protein Information Resource, University of Delaware); Ian Fore (National Cancer Institute, National Institutes of Health, Bethesda MD); Christian Haselgrove (University of Massachusetts Medical School); John Kunze (California Digital Library); Neil McKenna (Baylor College of Medicine); Pete Meyer, Harvard Medical School; Raman Prasad (IQSS, Harvard University); Peter Rose (University of California San Diego); Simone Sacchi, Columbia University); Ryan Scherle (Dryad Digital Repository); Curtis Smith (EndNote, Thomson Reuters); Cathy Wu (Protein Information Resource, University of Delaware).

Work on this project was coordinated by FORCE11 (https://force11.org), a not-for-profit community organization seeking to improve scholarly communication through digital technology.

Finally, the authors wish to thank Stephanie Hagstrom of the University of California Library and FORCE11 for her extremely helpful administrative work supporting the Data Citation Implementation Pilot, in organizing workshop and conference calls, and in coordinating website administration as well as logistics for workshop attendees.

